# 7 Tesla MRS estimates of GABA concentration relate to physiological measures of tonic inhibition in the human motor cortex

**DOI:** 10.1101/2024.07.12.603209

**Authors:** Ilenia Paparella, Paolo Cardone, Benedetta Zanichelli, Laurent Lamalle, Fabienne Collette, Siya Sherif, Mikhail Zubkov, William T Clarke, Charlotte J Stagg, Pierre Maquet, Gilles Vandewalle

**Affiliations:** GIGA-Research, CRC- Human Imaging Unit, 8 allée du Six Août, Batiment B30, University of Liège, 4000 Liège, Belgium; GIGA-Research, Coma Science Group, GIGA-Consciousness, University of Liège, Liège, Belgium; Wellcome Centre for Integrative Neuroimaging, FMRIB, Nuffield Department of Clinical Neurosciences, University of Oxford; Medical Research Council Brain Network Dynamics Unit, University of Oxford, Oxford, UK; Department of Neurology, Domaine Universitaire du Sart Tilman, Bâtiment B35, CHU de Liège, 4000 Liège, Belgium

**Keywords:** GABA, Magnetic resonance spectroscopy (MRS), TMS-EEG, neural mass model, inhibition, motor cortex

## Abstract

GABAergic neurotransmission within the cortex plays a key role in learning and is altered in several brain diseases. Quantification of bulk GABA in the human brain is typically obtained by Magnetic Resonance Spectroscopy (MRS). However, the interpretation of MRS-GABA is still debated. A recent mathematical simulation contends that MRS detects extrasynaptic GABA, mediating tonic inhibition. Nevertheless, no empirical data have yet confirmed this hypothesis. Here we collected ultra-high field 7 Tesla MRS and Transcranial Magnetic Stimulation coupled with high-density Electroencephalography (TMS-hdEEG) from the motor cortex of twenty healthy participants (age 23.95±6.4), while they were at rest. We first applied a Neural Mass Model to TMS-evoked potentials to disentangle the contribution of different GABAergic pools. We then assessed to which of these different pools MRS-GABA was related to by means of Parametric Empirical Bayesian (PEB) analysis. We found that MRS-GABA was mostly positively related to the NMM-derived measures of tonic inhibition and overall functionality of the GABAergic synapse. This relationship was reliable enough to predict MRS-GABA from NMM-GABA. These findings clarify the mesoscopic underpinnings of GABA levels measured by MRS and will contribute to the concretization of MRS-GABA promises to improve our understanding of human behaviour, brain physiology and pathophysiology.

**Key points:**

1. GABA neurotransmission is essential for synaptic plasticity and learning (especially motor learning) and is altered in several brain disorders, such as epilepsy and stroke.
2. Quantification of GABA in the human brain is typically obtained by Magnetic Resonance Spectroscopy (MRS). However, the interpretation of MRS-GABA is still debated.
3. By using a biophysical Neural Mass Model, here we show that MRS-GABA relates to physiological measures of tonic inhibition in the human cortex.

## Introduction

Gamma-aminobutyric acid (GABA) plays a crucial role in maintaining the excitation/inhibition balance in the brain, which is pivotal for optimal brain functioning^1^. GABA neurotransmission is essential for synaptic plasticity^2^ and learning (especially motor learning^3,4,5^) and is altered in several brain disorders, such as epilepsy and stroke^6^. Despite its functional significance, the interpretation of non-invasive GABA measurements in humans are challenged by the complex GABA signalling. After its release in the synaptic cleft, GABA binds with two major postsynaptic receptors, GABA_A_ and GABA_B_, which account approximately for 20% of the synapses in the cerebral cortex, hippocampus, thalamus, and cerebellum^7^. GABA is also found in the extracellular space where it binds to extrasynapatic GABA_A_ receptors^8^. These two different GABA pools play different roles in the brain with synaptic GABA triggering phasic inhibition and extracellular GABA mediating tonic inhibition^9^, which regulates neuronal gain and causes lasting modifications of neuronal activity, such as during learning.

Several techniques are available in humans to non-invasively probe specific aspects of GABA dynamics, ranging from molecular imaging (Magnetic Resonance Spectroscopy, MRS; Positron Emission Tomography, PET) to electrophysiology (Electroencephalography, EEG) and brain stimulation (Transcranial Magnetic Stimulation, TMS). None of these techniques accounts, alone, for all aspects of GABAergic signalling, making it therefore challenging to fully establish the contribution of phasic and tonic inhibition in human cognition. MRS allows for repeated, quantitative measurements of local GABA concentration without requiring invasive radio tracers (unlike PET) or a measurable motor output (unlike TMS), resulting in an enhanced clinical translatability. MRS is particularly promising when carried out at ultra-high-field MRI (≥7 Tesla, T), given the improved signal-to-noise-ratio and resolution compared to lower fields. Nevertheless, MRS-GABA cannot capture the dynamics of different GABAergic pools. A recent mathematical simulation suggested that MRS is most likely reflecting free extracellular GABA^10^ and could therefore provide a measure of tonic inhibition. However, this hypothesis is not yet supported by empirical data. Clarifying what is being measured by MRS would improve our understanding of MRS-GABA changes in behaviour and in neurological disorders, ultimately opening avenues for innovative care and treatment’s option for patients.

Here, we used a Neural Mass Model (NMM)^11^ of TMS-evoked brain potentials recorded with high-density EEG^12^, to model the dynamics of different GABAergic pools in a neuronal population of the motor cortex of 20 healthy subjects. We chose the motor cortex as motor evoked responses can be used as an objective means to standardise intensity and functional location of the TMS hotspot across participants and given GABA relevance for motor learning. NMM estimation of GABA activity has been validated by pharmacological interventions in humans and is considered equivalent to invasive interventions in animals^13,14,15,16^. We further collected ultra-high field 7T-MRS data over the same cortical area in the same healthy participants. We aimed to test whether MRS-GABA was associated to NMM’s estimation of tonic/phasic inhibition, with the hypothesis that GABA concentration would be related to physiological measures of tonic inhibition.

## Materials & Methods

### Participants

The study was approved by of the Ethical Committee of the CHU and Faculty of Medicine of Liège. Participants gave their written informed consent prior to their enrolment and received financial compensation for their participation. Exclusion criteria were as follows: BMI > 25 kg/m²; smoking or excessive alcohol consumers (> 14 units/week) or other addiction; use of sedative drugs, Na or Ca channel blockers, or other drugs acting on GABA/glutamate; metal inside the body, recent psychiatric history, severe head trauma, or sleep disorders, chronic medications; be left-handed. Twenty female participants were recruited (23.95y±6.44). Whether gender affects GABA concentration is still debated^17^. Some studies report contradictory gender differences in GABA (possibly due to regional variation)^18,19^, whereases other studies report no differences at all^20^, particularly in the motor cortex^21^. We decided to recruit only women to avoid potential sex biases and because mostly women responded to our initial call for participants. The hormonal phase of our participants was collected but was not used in the analyses. A strong bias arising from hormonal phase is unlikely in our design, given that TMS-EEG and MRS were collected on the same day or on consecutive days. Wake-up time and sleep duration/quality were also collected to avoid sleep deprivation from altering our results (see *Experimental Protocol*). **Table 1** shows all detailed participants’ demographics.

**Table 1.** Participants demographics. Characteristics of the total study sample. BMI: Body Mass Index. The hormonal phase is computed as the number of days from the start of the last menstrual cycle before participating in the study. Participants were also asked to communicate the start of the following menstrual cycle to take the length of their cycle into account. Wake-up time and scores of sleep duration/quality as reported in the self-completion Leeds Sleep Evaluation Questionnaire^22^. Values are provided as average ± standard deviation (SD).

|  | Sample (N=20) |
| --- | --- |
| **Age (y)** | 23.95y ±6.44 |
| **Sex** | Female |
| **BMI (kg/m<sup>2</sup>)** | 23.95 ± 3.87 |
| **Hormonal phase** | 40% Menstruation;<br>30% Follicular Phase;<br>25% Luteal Phase;<br>5% No period due to contraceptive pill |
| **Handedness** | Right |
| **Wake-up time** | 6h32min ± 20min |
| **Sleep duration** | 6h44min ± 1h |

### Experimental Protocol

Testing times were kept constant across participants to account for potential circadian variations in GABAergic activity^23^ (**Figure 1**). All participants came to the laboratory at 8 am. They were asked not to consume caffeine or alcohol containing beverages for >24h prior to their admission. They were asked to follow their normal sleep schedule the night before the study. Participants underwent a 30-minute structural scan at 7T MRI, which was later used for the TMS neuronavigation session and EEG source reconstruction. At around 9 am participants started the TMS-EEG session, which lasted for about 3h. After a small lunch break (caffeine/alcohol free), most participants (14) underwent an MRS session (at ∼3pm), which lasted around 1h. The voxel of interest (green box) was positioned over the same area previously stimulated with TMS-EEG by using anatomical landmarks. The remaining (6) participants decided to have the MRS on the following days (at 3pm) in which case the sleep requirement and questionnaire were repeated. For these participants we further ensured that the MRS session was kept at the same hormonal phase.

**Figure 1.**
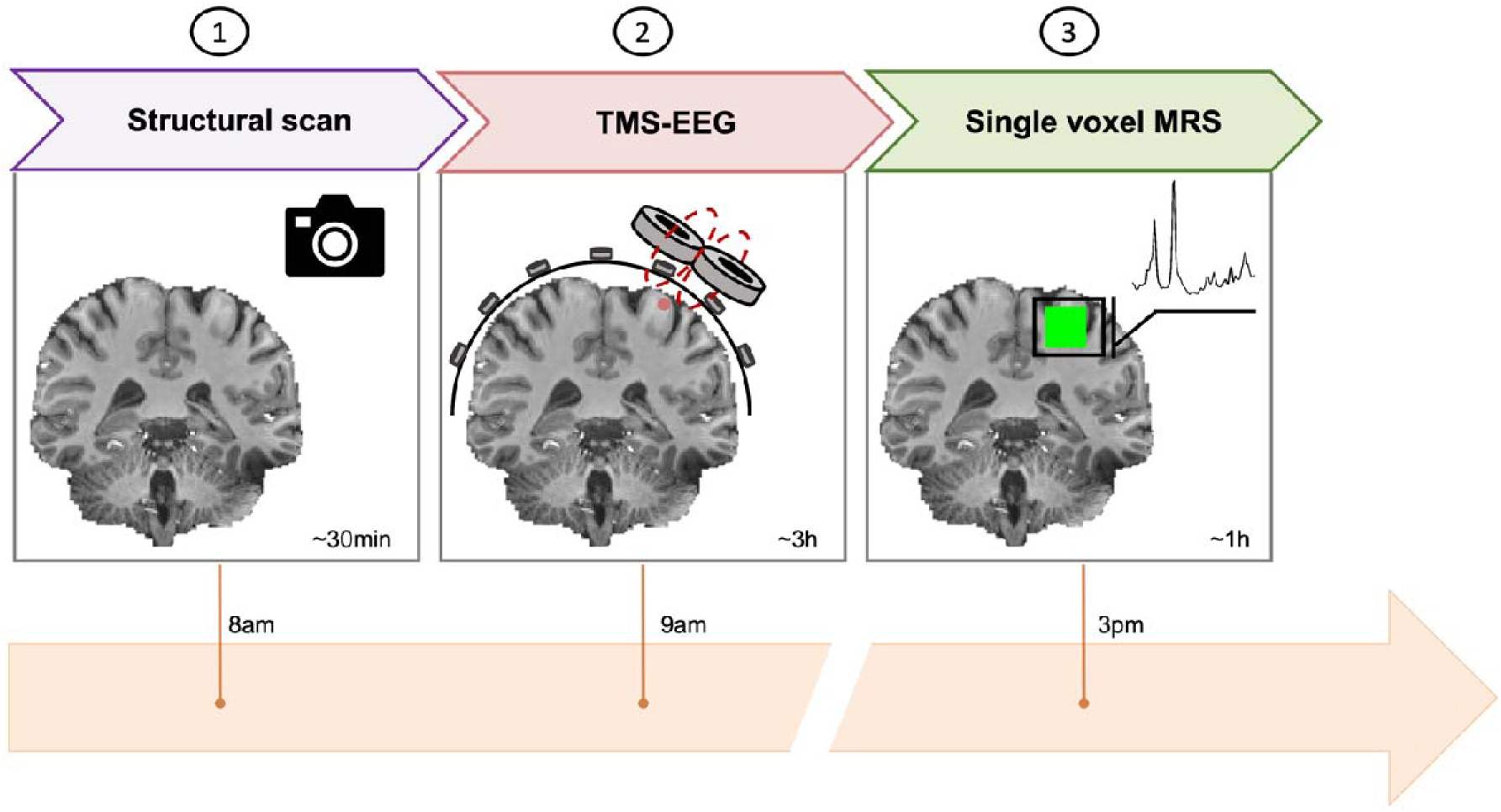
Graphical representation of the experimental protocol. All participants arrived at the lab at 8am. They first underwent a 30min structural scan at 7T MRI. At 9am, the TMS-EEG started. Our stimulation spot (here as red dot) was in the left motor cortex and was located on the structural scan. The TMS-EEG session lasted around 3h. Afterwards, participants underwent an MRS session at 7T MRI (roughly at 3pm). The voxel of interest (here represented as a green box) was positioned over the same area previously stimulated with TMS-EEG by using anatomical landmarks. The MRS session lasted around 1h. An example cropped spectra is also provided.

#### Data acquisition

##### Structural T1 scan

A T1-weighted structural image was acquired using a 7T MRI system (Terra, Siemens Healthineers, Erlangen, Germany) with a 32-channel receiver and 1-channel transmit head coil (Nova Medical, Wilmington, MA, USA). Structural MRI data were acquired with a Magnetization Prepared Rapid Acquisition Gradient Echo (MPRAGE) sequence: TR, 2300ms; TE, 2.76ms; slice thickness, 1.0mm; in-plane resolution 1.0 x 1.0 mm^2^, GRAPPA factor =2. The acquired structural image was used to inform the neuronavigation during the TMS-EEG session.

##### TMS-EEG

TMS-EEG data were recorded with a 60-channel TMS-compatible EEG amplifier (Eximia EEG, Nexstim, Helsinki, Finland), and TMS was delivered by means of a Focal Bipulse 8-Coil (Eximia TMS) combined with a magnetic resonance–guided navigation system (eXimia NBS). The EEG amplifier gates the TMS artefact and prevents saturation by means of a proprietary sample-and-hold circuit that ensure the absence of TMS-induced magnetic artefacts from 8ms post-TMS.

Our stimulation site was the right motor hotspot for the first dorsal interosseous (FDI) on the left hemisphere, defined as the scalp location where the largest and most consistent MEPs could be identified in the FDI. The TMS coil orientation was then adjusted to obtain an artifact-free TMS-evoked potential (TEP) following recent recommendations^24^. Resting motor threshold was defined as the minimum intensity needed to elicit a peak-to-peak motor evoked response with an amplitude of more than 50μV in at least 5 out of 10 subsequent trials while the targeted muscle was at rest.

The stimulation session consisted of 250 pulses at a stimulation intensity below resting motor threshold to reduce muscular contamination in the EEG^25^. Electromyography (Nexstim EMG) was recorded via disposable ECG electrodes placed over the FDI of the right hand, using a belly-tendon montage with a ground electrode over the ulnar styloid process. During the session, participants were seated on a comfortable reclining bed, with their eyes open and their right hand positioned on a pillow placed over their lap. They were instructed to fixate a black dot placed on the wall in front of them at eye level and to keep the targeted muscle relaxed while EMG was continuously monitored on a computer screen. The Inter Stimulus Interval (ISI) was randomly set to 1900–2200ms.

EEG signal was band-pass filtered between 0.1 and 500Hz and sampled at 1450 Hz. All electrode impedances were maintained <5kΩ. Electro-oculogram was recorded with two additional bipolar electrodes. During the EEG recording, participants’ perception of the clicks produced by the TMS coil discharge was eliminated using earplugs continuously playing a pink masking noise (< 90dB, adjusted per participant prior to each recording). Bone conductance was minimized by applying a thin foam layer between the EEG cap and the TMS coil. The absence of an auditory evoked response was verified during each recording and was confirmed by delivering, in a short sham session, 30-40 pulses parallel to the scalp while the masking noise was played at the same level. At the end of each recording, electrode positions on the participants’ head were recorded. Participants were continuously monitored by a member of the team to ensure they did not fall asleep or doze off during the session.

#### MRS

MRS data were acquired with the same 7T MRI system. Participants were instructed to lay still at rest while in the MRI scanner without falling asleep. Dielectric pads (Multiwave Imaging, Marseille, France) were placed over the left central sulcus to increase B1 efficiency over the M1 voxel of interest (VOI). B1 efficiency was imaged using actual flip angle imaging (AFI): field of view 240 x 240 mm^2^, repetition times (TR1/2) 6 / 30 ms, Echo Time (TE) 2.58 ms; non-selective flip angle 60°; slice thickness 2.5 mm, in-plane resolution. To enable the placement of the MRS voxel, we repeated the T1 structural acquisition as detailed above. Coronal and axial images were resampled from the sagittal T1-weighted image and used to place a 2 x 2 x 2 cm^3^ VOI. Anatomical landmark from the screenshot of the motor hotspot stimulated in the previous TMS-EEG session were used to guide voxel positioning over the T1 image. **Figure 2** provides an heatmap of the VOI and TMS-EEG spot across all participants. Shimming was performed in two steps: first using the vendor’s own GRE shimming, second using FASTMAP^26^. MRS data were acquired using a semi-LASER sequence (localisation by adiabatic selective refocusing) provided as part of the Center for Magnetic Resonance Research, (University of Minnesota) Spectroscopy package SEMI-LASER (MRM 2011, NMB 2019) Release 2016-12, 10.1002/nbm.4218: TR = 5000 ms, TE = 26 ms (TE1/2/3 = 7 ms, 10 ms, 9 ms), 20 x 20 x 20 mm^3^ voxel, 64 averages per block, TA = 5min 20s, using per-subject calibrated VAPOR (Variable Power RF pulses with Optimized Relaxation delays) water suppression. Further details on the sequence are included in the *MRSinMRS* checklist we provide as **Supplemental Information**.

**Figure 2.**
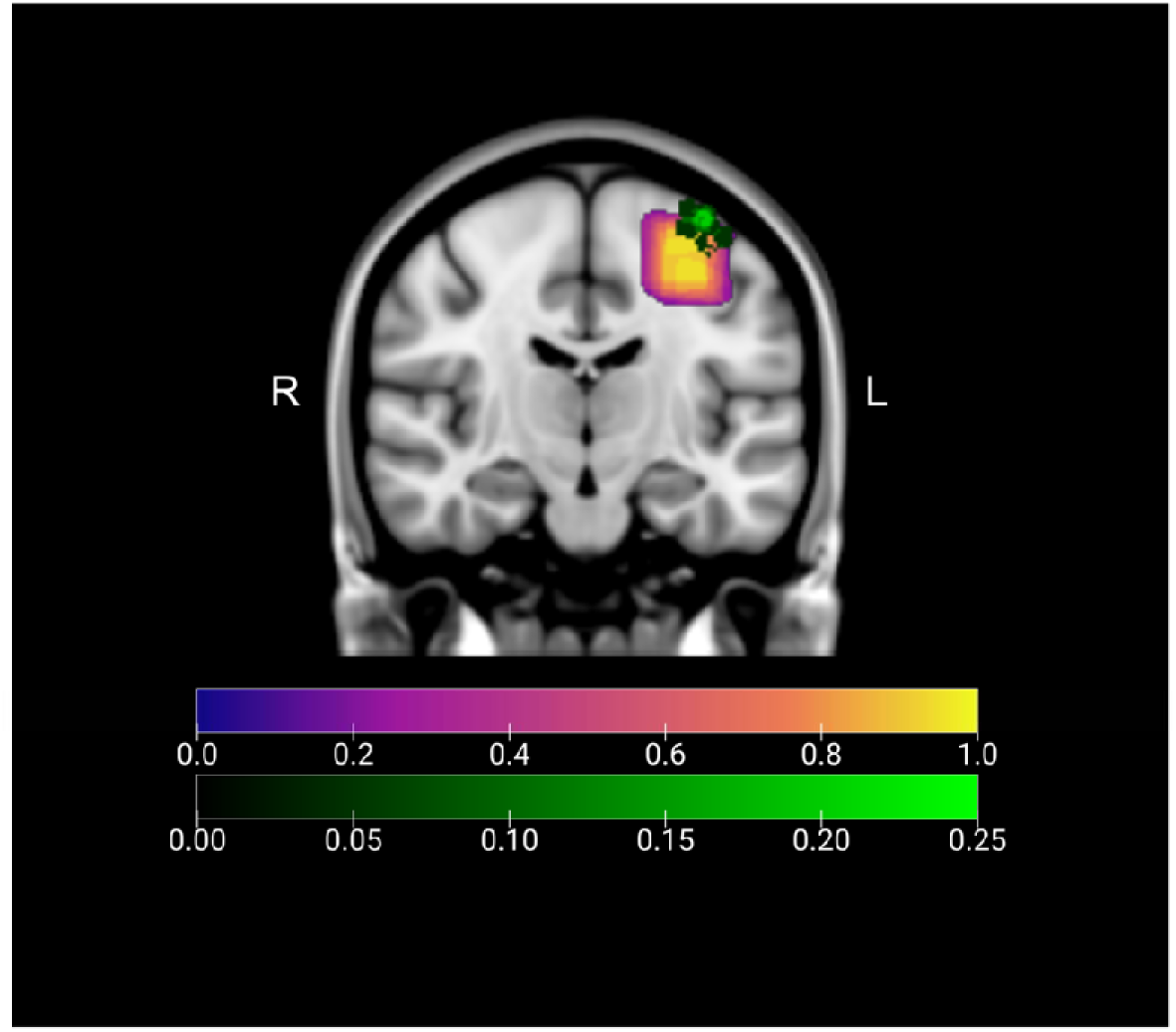
Probabilistic heatmap of MRS VOI placement and TMS-EEG hotspot location across participants. The heatmaps are shown on the coronal view of the MNI_152_T1_1mm template (coordinates [−37, −18, 64]) provided by FSL^27^. The colour scales represent the proportion of participants whose VOI (upper) or TMS-EEG spot (lower) overlapped in that voxel. As a rule, each VOI was placed in the left primary motor cortex, as close as possible to the TMS-EEG spot (by following anatomical landmarks), and avoiding non-brain tissue (e.g., the dura) that could induce artifacts in the signal.

#### Data analysis

##### TMS-EEG & NMM

TMS-EEG data were preprocessed with Statistical Parametric Mapping 12 (SPM12, http://www.fil.ion.ucl.ac.uk/spm/) and analysed with MATLAB R2019 (MathWorks, Natick, Massachusetts). Data were visually inspected to reject artifacted channels and trials (blink, body movements, slow eye movements and other artifacts; rejection rate of around 10%). EEG signals were re-referenced to the average of all good channels. Continuous EEG recordings were lowpass-filtered at 80 Hz, down-sampled from 1450 to 1000 Hz and then high-pass-filtered at 1 Hz. Individual trials were then epoched between −100 pre and 300ms post-TMS pulses. Baseline correction (−100 and −1.5ms pre TMS pulses) was applied. Robust averaging was used to compute the mean evoked response.

We then used NMM^11^, within the Dynamic Causal Modelling (DCM)^28^ framework, to model our evoked response. NMMs, originally used to study multi-region network responses, are applied more and more to describe molecular factors underlying brain activity in a single region. NMM models event-related potentials as the response of a network to exogenous inputs, by inferring neuronal states within a given cortical area comprising 4 subpopulations of neurons, namely deep and superficial pyramidal cells, excitatory stellate cells, and inhibitory interneurons. These subpopulations exhibit self-inhibition controlling neuronal gain and communicate through excitatory and inhibitory connections^11^. The model also include active currents that describe ligand-gated excitatory and inhibitory ion flow, mediated through glutamatergic (α -amino-3-hydroxy-5-methyl-4-iso-xazolepropionic acid receptors, AMPA, and N-methyl-D-aspartate, NMDA) and GABAergic (GABA_A_) receptors^13^. GABA_A_, AMPA, and NMDA measures are the rate constant of the receptors lumped over the entire circuit. A graphical representation of the NMM is provided in **Figure 3A**.

**Figure 3.**
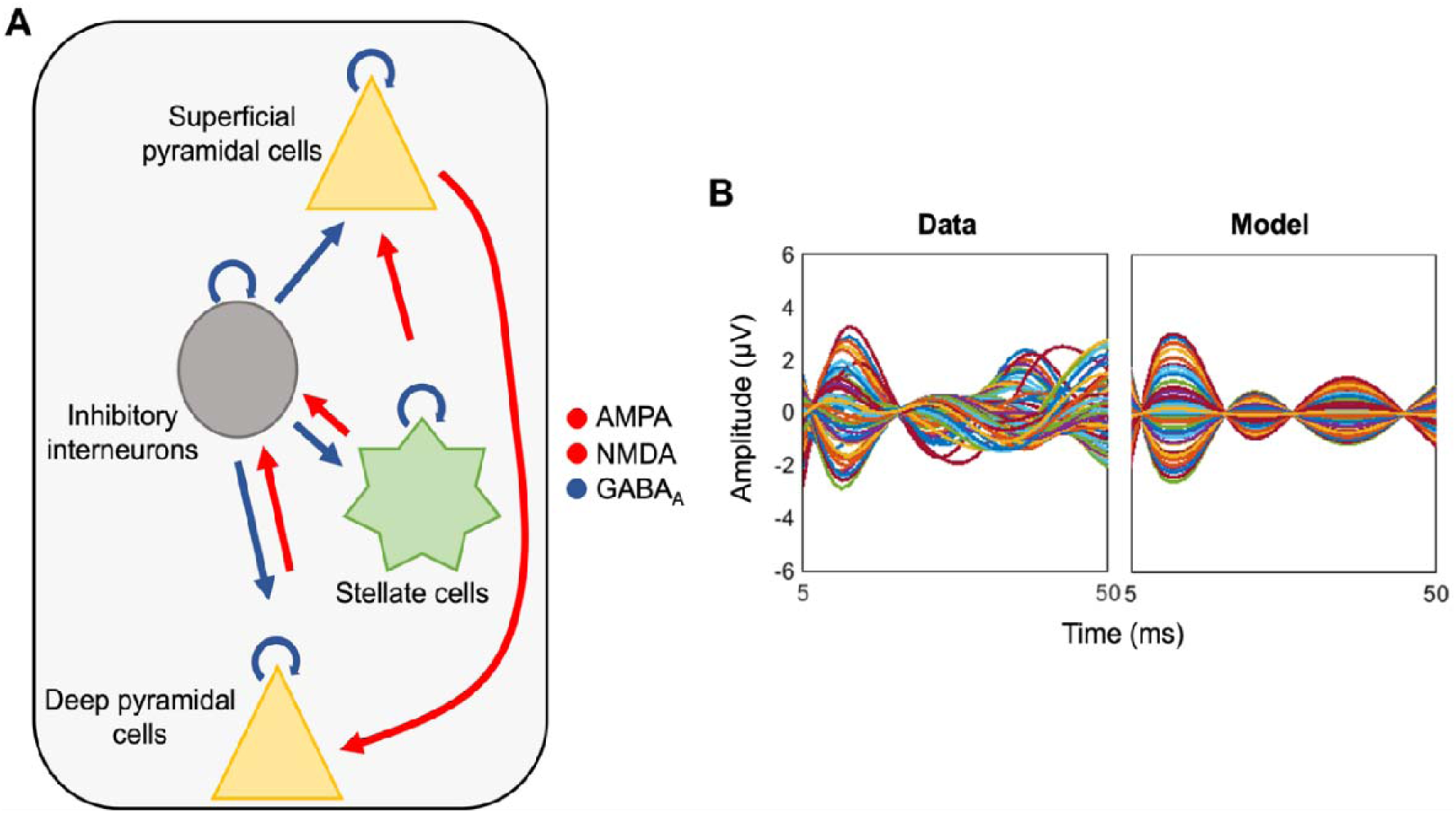
Neural Mass Model and example of model fits. **A.** NMM decomposes a cortical area into 4 neuronal subpopulations: superficial and deep pyramidal cells (yellow), spiny stellate cells (green) and inhibitory interneurons (grey). Each subpopulation is assigned to a particular cortical layer and projects to the other subpopulations via anatomically plausible excitatory (red) and inhibitory (blue arrows) connections and have inhibitory feedback-loops controlling neuronal gain. NMM also provides information on the density and the activity of three mains receptors: AMPA and NMDA (red), mediating excitatory connections and GABA_A_ mediating inhibitory ones (blue). This figure was simplified from Paparella et al., 2023^6^. **B.** Example of model fit with an observed (left) and modelled (right) TMS evoked response for one of our participants. The response is plotted as a function of time and amplitude. Each line represents a channel.

Importantly, this model allows for the distinction between tonic background inhibition, mediated by cell inhibitory self-connections, and phasic inhibition, modelled by the projections from interneurons to the other cells^29^.

The TMS pulse is assumed to represent the input as would be typically induced by sensory stimuli. To estimate which NMM parameter (and how) contributes to the evoked response, DCM uses generative or forward models for evoked EEG responses and fits these models using a variational Bayesian inversion scheme. We modelled the active source (individual MRI coordinates of the TMS hotspot) by means of a single equivalent current dipole (ECD) within an electromagnetic forward model over the 5–50ms TEP, a time window where changes in EEG can arguably be attributed to the sole local effects of the TMS. This model used a ‘boundary element method’ approach, with homogeneously and isotropically conductive volumes delimited by the brain, cerebrospinal fluid (CSF), skull and scalp surfaces. Individual head models are derived using an inverse spatial normalization of a canonical mesh for each participant (MRI T1-sequence, 20400 dipoles). The position of the 60 electrodes was coregistered in each participant before forward model computation. A lead-field mapping of cortical sources onto measured signals was parameterized for orientation and location of the ECD. In the NMM model we used, all the parameters were allowed to vary, to enable the model to recreate complex neurophysiological brain states. An example of observed and modelled data is provided in **Figure 3B**.

##### MRS

Quantitative analyses of the spectra were performed using the open source FSL-MRS^30^ analysis toolbox. Raw (“twix”/“.dat”) MRS data were first converted into the standard data format for magnetic resonance spectroscopy (NIfTI-MRS) using the conversion program *spec2nii*^31^. Converted data were then pre-processed with *fsl_mrs_preproc,* the pre-packaged processing pipeline for non-edited single voxel MRS data provided by FSL-MRS. The pipeline performs all the processing operations recommended in the community-driven consensus paper^32^, including coil combination, phase and frequency alignment of individual transients, averaging, eddy current correction, removal of residual water with HLSVD, and zero-order phase correction. *Fsl_mrs_sim* was used to create a set of basis spectra by providing precise description of the sLASER sequence (timings, RF pulses, and relevant gradients). An empirically measured macromolecular baseline was added to the basis set. The generated basis spectra were then fitted to the data using a Markov chain Monte Carlo (MCMC) optimization. To quantify the proportion of white matter, grey matter and CSF in the VOI, the *svs_segment* command was applied to the T1 image, previously preprocessed using the fsl_anat structural processing pipeline in the FMRIB software library (FSL)^27^, which performs some pre-processing steps including bias correction, brain extraction^33^, registration to standard space^34^ and tissue segmentation^35^. All the metabolites were corrected for the proportion of total brain tissue in the VOI. There was no strong correlation (> ±0.3) between GABA and other metabolites, indicating good spectral separation was achieved. The exclusion criteria for the data were as follows: water linewidths at full width at half maximum (FWHM) > 15 Hz or signal/noise ratio (SNR) < 30^36^. The quality check was performed through the *fsl_mrs_summarise* command. None of the data met the exclusion criteria, thus all were included for further analysis. GABA concentration is expressed as a ratio of total Creatine (tCR - validated reference compound in GABA MRS^37^) and reported as MRS-GABA. Further details on the analysis steps are included in the *MRSinMRS* checklist we provide as Supplemental Information

### Statistics

NMMs describes the average microscopic activity of the neural subpopulations within a cortical column in the region of interest. The first-level analysis infers neural responses of individual TMS-EEG data by inverting the Bayesian model^38,39^, which allows finding the parameters that offer the best trade-off between predictions and observations. Once we estimated the NMM for each subject, we then tested whether MRS-GABA was related to any of the parameters in our model using Parametrical Empirical Bayes (PEB)^40^ as included in SPM12. To avoid dilution of evidence, we reduced the search space^40^ by carrying out a PEB analysis only over the T and H matrix of the NMM. These two matrices represent receptor densities (NMDA, AMPA and GABA_A_) and excitatory/inhibitory connections among sub- populations of neurons, respectively, and they were selected as they are the only parameters in the NMM providing GABAergic information both in isolation and in interaction with the glutamatergic counterpart. In PEB, the probability densities of all parameters of the NMM are collated and modelled at the second level with any unexplained between-subject variability captured by a covariance component mode. PEB then performs Bayesian Model Reduction (BMR), which ‘prunes’ any parameter not contributing to the model evidence by automatically searching over reduced models (those with some parameters switched off). Then, Bayesian Model Average (BMA) is calculated over the models from the final iteration of the greedy search. Only NMM parameters with a strong significant relationship with MRS-GABA, as indexed by a posterior probability (Pp) > 0.99, are reported in the results section. To provide visual representation of our results, individual peak estimates were extracted to obtain linear regression plots. We then used leave-one-out (LOO) to test the meaningfulness of the relationship (predictive validity) found between MRS- and NMM-GABA. A PEB model was fitted to all but one subject, and covariates for the left-out subject were predicted. This was repeated with each subject left out and the accuracy of the prediction was recorded. Optimal sensitivity and power analysis in DCM/PEB remains under investigation. We nevertheless computed a prior sensitivity analysis to get an indicator of the minimum detectable effect size in our main analyses given our sample size. According to G*Power 3 (version 3.1.9.4)^41^ taking into account a power of .8, an error rate of .05, a sample size of 20 allowed us to detect a medium effect sizes r>.27 (confidence interval: −0.2, 0.64; R²>.07, R² confidence interval: .04-.41) within a linear multiple regression framework including 1 predictor. Based on this, we deemed the sensitivity reasonable.

## Results

The PEB analysis revealed that MRS-GABA correlated positively with the self-inhibitory feedback loop of almost all the subpopulation of cells included in the model, namely the inhibitory interneurons, stellate and deep pyramidal cells (**Figure 4A**). PEB further showed that MRS-GABA was negatively related with the excitatory connection from stellate to superficial pyramidal cells and to the inhibitory connection from inhibitory interneurons to deep pyramidal cells. At the receptors level, the only significant effect was a positive relationship with the activity of GABA_A_. The effect sizes of each significant relationship are reported in **Figure 4A** and represent the rate of change of the NMM parameter as a function of MRS-GABA, with the sign specifying the direction of the relationship. As an example, the effect size of the relationship between MRS-GABA and the self-inhibitory feedback loop of the inhibitory interneuron is 5.15, meaning that when MRS-GABA increases one fold, the inhibition of the inhibitory interneurons increases as well by 5.15 times (in the units of the underlying DCM). **Figure 4** includes two linear regression plots to represent the positive relationship found between MRS-GABA and NMM-tonic inhibition (calculated as the mean of the significant self-inhibitory feedback loops’ peak estimates) (**Figure 4B**) and between MRS-GABA and the NMM-GABA_A_ receptor activity (**Figure 4C**). These plots are for display purposes only and do not substitute the outcomes of the PEB analysis.

**Figure 4.**
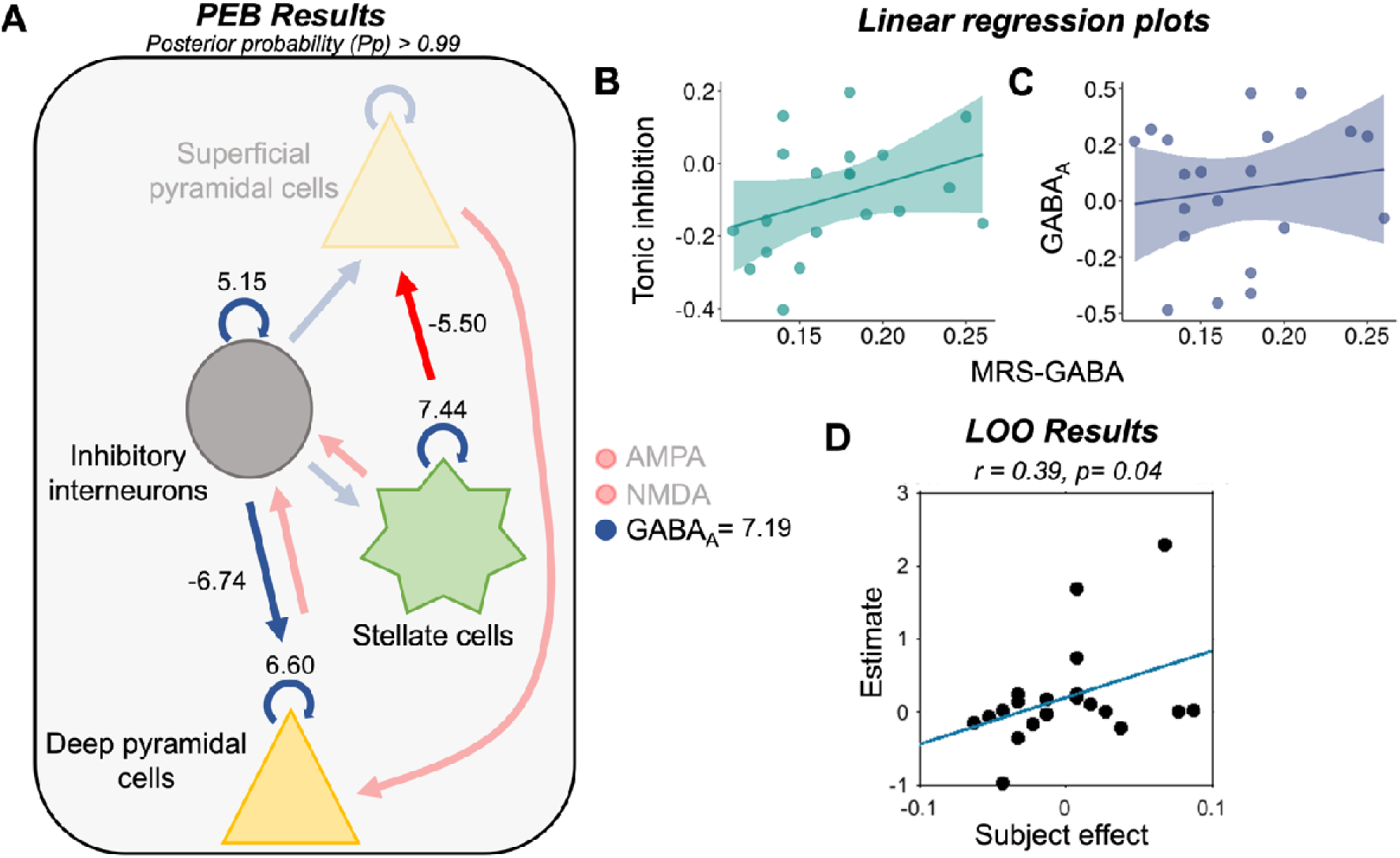
Association between NMM parameters and MRS GABA. **A.** Outcomes of the PEB analyses. Only parameters showing a significant relationship with MRS-GABA (Pp>0.99) are highlighted. The sign indicates the direction of the relationship (positive or negative) whereas the values indicate the rate of change of each parameter as a function of MRS-GABA. Overall, we observed that MRS-GABA was positively related to the tonic inhibition of the inhibitory interneurons, stellate cells and deep pyramidal cells, whereases was negatively related to the excitatory connection from stellate to superficial pyramidal cells and to the inhibitory connection from inhibitory interneurons to deep pyramidal cells. Among the receptors, MRS-GABA was shown to be only positively related to the activity of GABA_A_. **B&C.** Regression plots representing the positive effects found between MRS-GABA and the NMM-derived GABAergic measures: tonic inhibition, calculated as the mean of the significant self-inhibitory feedback loops’ peak estimates, (B) and the activity of the GABA_A_ receptor (C). These plots are for display purposes only and do not substitute the outcomes of the PEB analysis. **D.** LOO analysis. The actual subject effect is plotted against the expected value of the estimated subject effect. This analysis allows to test whether the positive relationship found between NMM-self_II and MRS-GABA is meaningful enough that one could predict one based to the other. We found a significant relationship between the actual MRS-GABA and the predicted MRS-GABA for each left-out subject (Pearsons’s rho=0.39, p=0.04).

We then wanted to further assess whether the size of the positive relationship between NMM-tonic inhibition and MRS-GABA was meaningful enough that we could predict one with the other. Assessing predictive validity is particularly important for studies determining the clinical significance of model parameters. Several NMM parameters showed a significant relationship with MRS-GABA, and we could not use them all for the predictive analysis, especially given our small sample size. We used the NMM parameter encoding for the self-inhibitory feedback loop of the inhibitory interneurons (NMM-self_II) for our next analysis as it constitutes the most canonical proxy of tonic inhibitory activity within the NMM. We then used LOO to test if we could predict MRS-GABA from NMM-self_II. The out-of-samples correlation of the actual MRS-GABA over the (expected value of) the predicted MRS-GABA for each left-out subject was significant (Pearsons’s rho=0.39, p=0.04) (**Figure 4D**). Therefore, the relationship between NMM-self_II and MRS-GABA was sufficiently large to predict the left-out subjects’ MRS-GABA with performance above chance.

## Discussion

MRS enables non-invasive *in vivo* measurements of neurotransmitter concentration in the brain. Despite the increasing number of MRS-based findings on the role of GABA in human behavior, learning and neurological disorders, their relevance in terms of cortical networks dynamics remains uncertain. This uncertainty limits the interpretation of MRS results, especially when one wants to relate MRS-GABA to behavior or pathological processes.

Here we collected 7T MRS and TMS-EEG data over M1 in twenty young healthy participants. We then applied a biophysical NMM on our electrophysiological data to establish potential relationships between modelled neurophysiological parameters and MRS. We demonstrated that MRS-GABA was related most closely to tonic inhibition, being positively associated with the interneurons, stellate cells, and deep pyramidal cells’ inhibitory feedback loops. MRS-GABA was also associated with the functioning of the GABAergic synapse at large, as indexed by the GABA_A_ parameter, which mediates all inhibitory connections allowed in the model. The relationship observed between MRS- and tonic NMM- GABA was meaningful enough that we could predict the left-out subjects’ MRS-GABA from the NMM-GABA, taking as proxy of NMM tonic inhibition the activity of the inhibitory interneurons’ feedback loop.

### NMM helps clarifying what is being measured by MRS-GABA

The morphology of the motor cortex^42,43^ is now well established. The majority of neurons are pyramidal cells, although 28% are interneurons, of which the majority are excitatory or spiny stellate cells^44^. Stellate cells are mostly found in layer 4 of the cortex^45^, and for many years it was thought that the motor cortex did not have a layer IV, suggesting that the neural circuitry controlling motor movements was different from other cortical regions. However, multiple lines of evidence have challenged this view by showing that neurons at the border between layer 3 and 5 of the motor cortex possess many of the same proprieties of stellate cells^46^ and are distributed in several layers, even in more superficial ones^47^. The convoluted and fine-tuned connections between pyramidal cells and interneurons are used to adjust the balance between excitation and inhibition and finely select the output of the circuit^48^.

To best capture the functioning of a cortical motor network, the NMM used in this study comprised all three abovementioned cell types with the pyramidal cells further subdivided into superficial and deep. The resulting four subpopulations of neurons, whose activity is controlled and regulated by self-inhibitory feedback loops, form excitatory or inhibitory connections mediated by AMPA and NMDA or GABA_A_ receptors, respectively. TMS is thought to enter this network by inducing a strong depolarization of the superficial pyramidal cells^49^, which, in turn, leads to the recruitment of fully synchronized clusters of deep pyramidal neurons and interneurons to control the firing of excitatory networks and adjust the E/I balance^48^. To further describe the complex cortical circuitry, our NMM distinguishes between inhibitory self-connections and inhibitory projections from interneurons to the other cells. Self-connections are assumed to mediate tonic background inhibition, while intrinsic inputs to pyramidal and stellate cells can be regarded as mediating phasic inhibition^12^.

When using PEB analysis to reveal potential associations between MRS-GABA and NMM parameters we observed that MRS-GABA was positively related to the self-inhibitory feedback loops (i.e., tonic inhibition) of almost all neural subpopulations, but the superficial pyramidal cells, which were most probably receiving the TMS input. Higher MRS-GABA was associated to higher tonic inhibition of deep pyramidal cells, stellate cells, and inhibitory interneurons. Consequently, more inhibited stellate cells excited less superficial pyramidal cells whereas more inhibited interneurons inhibited less deep pyramidal cells. The analysis also revealed a positive relationship between MRS-GABA and GABA_A_ receptor activity, which is a time constant that encodes how fast inhibitory post-synaptic potentials recover after a presynaptic spike has reached the dendrite, thus reflecting the functioning of the inhibitory synapse at large.

Our findings align with several other studies that tried to relate MRS-GABA to measures of phasic inhibition in humans and found no significant relationships. Phasic inhibition was either assessed through TMS using paired pulses with different interstimulus intervals^50–53^, or through [^11^C]flumazenil PET, a radiotracer that binds to the benzodiazepine site of GABA_A_ receptors thus providing a more direct quantification of the receptors availability at the level of the synapse^54^. MRS-GABA showed only one significant relationship with what is believed to be a paired-pulses TMS measure of tonic inhibition^50,55^, even though this result has not been replicated in more recent studies^52,53^. By showing that MRS-GABA mostly relates to parameters that encode for tonic inhibition in an biophysical NMM, our study confirms empirically what a recent mathematical simulation seemed to suggest^10^, and may further explain why the previous attempts to relate MRS-GABA to measures of phasic inhibition, failed.

Our study does not come without flaws. First, the sample size is relatively small. A higher sample size would have increased statistical power, though relationships with meaningful effect sizes should be evident with this sample size. Second, our sample comprised females only, limiting the generalizability of the findings. Finally, and most obvious, NMM does not provide a direct measure of the actual biological process but a validated approximation. Invasive electrophysiological measurements in pathological brains of patients undergoing surgery could partly address this limitation although interneurons are more difficult to record using multiunit recordings. Future studies could also assess MRS-GABA before and after pharmacological manipulation of ambient/tonic GABA. Likewise, animal models could combine ultra-high field MRS at 9.4T ^104^ and opto- and chemogenetics to further validate the mechanistic links between MRS signal and GABAergic neurotransmission^56^.

### The importance of elucidating GABAergic measures in the clinical context

GABAergic inhibition has a major impact on information processing, plasticity and network synchronization^57^, and its clinical relevance in many neurological disorders has been proven. Epilepsy is an obvious example. Changes in inhibitory neurotransmissions are found both in animal models^58^ and human epileptic patients^59^. Epileptogenic phenomena have been associated with episodes of excitatory or inhibitory cellular hyperactivity, changes in traffic and expression of GABA receptors and disequilibrium between tonic inhibition and neuronal excitability in animals^60^. In humans, studies testing GABAergic changes in epilepsy showed increased GABA levels in patients with juvenile myoclonic epilepsy^61^, idiopathic generalized epilepsy^62^ and refractory focal epilepsy^63^, which were associated with malformations of cortical development compared to controls.

Beyond epilepsy, stroke represents another clinical example of disruption and imbalance between excitatory and inhibitory activity in the brain. Studies investigating GABAergic alterations post stroke, revealed a lower GABA concentration in the contralesional M1^64^, decreased GABA concentration following constraint-induced motor therapy^65^, and lower GABA_A_ receptor activity in the ipsilesional M1^66^, causing an overall higher excitation/inhibition ratio in the affected hemisphere^67^, which was inversely correlated with motor recovery^68^.

GABAergic changes have been highlighted in other neurological and psychiatric disorders, such as relapsing-remitting multiple sclerosis^69^, Alzheimer’s^70^ and Parkinson’s^71^ disease, substance use disorder^72^, schizophrenia^73^, and autism spectrum disorder^74^, to name a few. Most of the studies bringing to these results use MRS to quickly and non-invasively quantify GABA in patients. Knowing that these GABAergic alterations found in patients versus controls do not apply to any type of GABA but specifically to tonic GABA, may inform novel interventions and therapeutic targets to ensure more tailored treatments. The development of drugs able to alter tonic GABA by acting on extrasynaptic GABA receptors has seen remarkable progress. For instance, GABAmimetic drugs, such as muscimol and THIP (gaboxadol; 4,5,6,7-tetrahydroisoxazolol[4,5-c]pyridine-3-ol), have been shown to act on GABA_A_ receptor agonist sites with preferential activation of the high-affinity extrasynaptic receptors^75^.

## Conclusions

MRS is increasingly being used to investigate GABAergic changes in inter-individual behavioural differences and in the context of neurological disorders. Recent advances in MRS, such as high-resolution 3-D imaging of metabolic profiles over large brain regions by Magnetic Resonance Spectroscopy Imaging (MRSI) at ultra-high field^76^, increase the promise of MRS as clinical tool. Our study provides empirical data to clarify the mesoscopic underpinnings of MRS-GABA, supporting the idea that it primary measures extrasynaptic GABA activity (i.e., tonic inhibition). Elucidating its functional significance will improve our understanding of human behaviour, brain physiology and pathophysiology.

## Acknowledgements

The authors thank Daniel Hauke and Christophe Phillips for valuable discussions over Neural Mass Models and his help in the interpretation of the model output. We also thank Annick Claes, Christian Degueldre, Brigitte Herbillon, Patrick Hawotte, Benjamin Lauricella, Eric Salmon, and André Luxen for their help over the different steps of the study.

## Author contributions

I.P., G.V., and P.M. designed the research. I.P. and P.C., set up the experiment. I.P., P.C., and B.Z. acquired the data. C.J.S., W.C., and S.S. supported data acquisition by providing MRS sequences. L.L., F.C., and M.Z. gave important insights while discussing and interpreting the data. I.P. analyzed the data supervised by G.V. regarding the TMS-EEG part, and by W.C. and C.J.S. for the MRS part. I.P., P.M. and G.V. wrote the paper. All authors edited and approved the final version of the manuscript.

## Data availability statement

The processed data and analysis scripts supporting the results included in this manuscript are publicly available via the following open repository: https://gitlab.uliege.be/CyclotronResearch-Centre/Public/fasst/XXX (created after acceptance). The raw data could be identified and linked to a single subject and represent a large amount of data. Researchers willing to access to the raw should send a request to the corresponding author (GV). Data sharing will require evaluation of the request by the local Research Ethics Board and the signature of a data transfer agreement (DTA).

