## Supplementary Materials for "7 Tesla MRS estimates of GABA concentration relate to physiological measures of tonic inhibition in the human motor cortex"

**Supplementary information**

*MRSinMRS checklist for a 7T MRS study*

In line with the need of minimum guidelines for the reporting of MRS methods and results^2^, we completed the MRSinMRS checklist where the standardized description of MRS hardware, data acquisition, analysis, and quality assessment is provided.

| 1. Hardware |  |
| --- | --- |
| a. Field strength [T] | 7 T |
| b. Manufacturer | Siemens Healthineers, Erlangen, Germany |
| c. Model (software version if available) | Magnetom 7T |
| d. RF coils: nuclei (transmit/receive), number of channels, type, body part | 32-channel receiver and 1-channel transmit head coil (Nova Medical, Wilmington, MA, USA) |
| e. Additional hardware | / |
| 2. Acquisition |  |
| a. Pulse sequence | Semi-LASER |
| b. Volume of interest and VOI locations | Single voxel placed over the hand knob located in the motor cortex |
| c. Nominal VOI size [cm^3^, mm^3^] | Anatomy-matched, 2x2x2 cm^3^ |
| d. Repetition Time (TR), Echo Time (TE) [ms, s] | / |
| e. Total number of excitations or acquisitions per spectrum (NA)  In time series for kinetic studies   i. Number of averaged spectra) per time-point (NA) ii. Averaging method (e.g. block-wise or moving average)   Total number of spectra (acquired / in time-series) | */* |
| f. Additional sequence parameters (spectral width in Hz, number of spectral points, frequency offsets)   i. If STEAM:, Mixing Time (TM) ii. If MRSI: 2D or 3D, FOV in all directions, matrix size, acceleration factors, sampling method | / |
| g. Water suppression method | / |
| h. Shimming method, reference peak, and thresholds for “acceptance of shim” chosen | */* |
| i. Triggering or motion correction method  (respiratory, peripheral, cardiac triggering, incl. device used and delays) | / |
| 3. Data analysis methods and outputs |  |
| a. Analysis software | Both processing and fitting was done in FSL-MRS^3^ |
| b. Processing steps deviating from quoted reference or product analysis software (vendor, version) | / |
| c. Output measure  (e.g. absolute concentration, institutional units, ratio) Processing steps deviating from quoted reference or product | / |
| d. Quantification references and assumptions, fitting model assumptions | GABA was quantified as referenced to total creatine |
| 5. Data Quality |  |
| a. Reported variables  (SNR, Linewidth (with reference peaks)) | / |
| b. Data exclusion criteria | The exclusion criteria for the data were as follows: water linewidths at full width at half maximum (FWHM) > 15 Hz or signal/noise ratio (SNR) < 30 |
| c. Quality measures of postprocessing Model fitting (e.g. CRLB, goodness of fit, SD of residual) | */* |
| d. Sample Spectrum | Figure S1 |

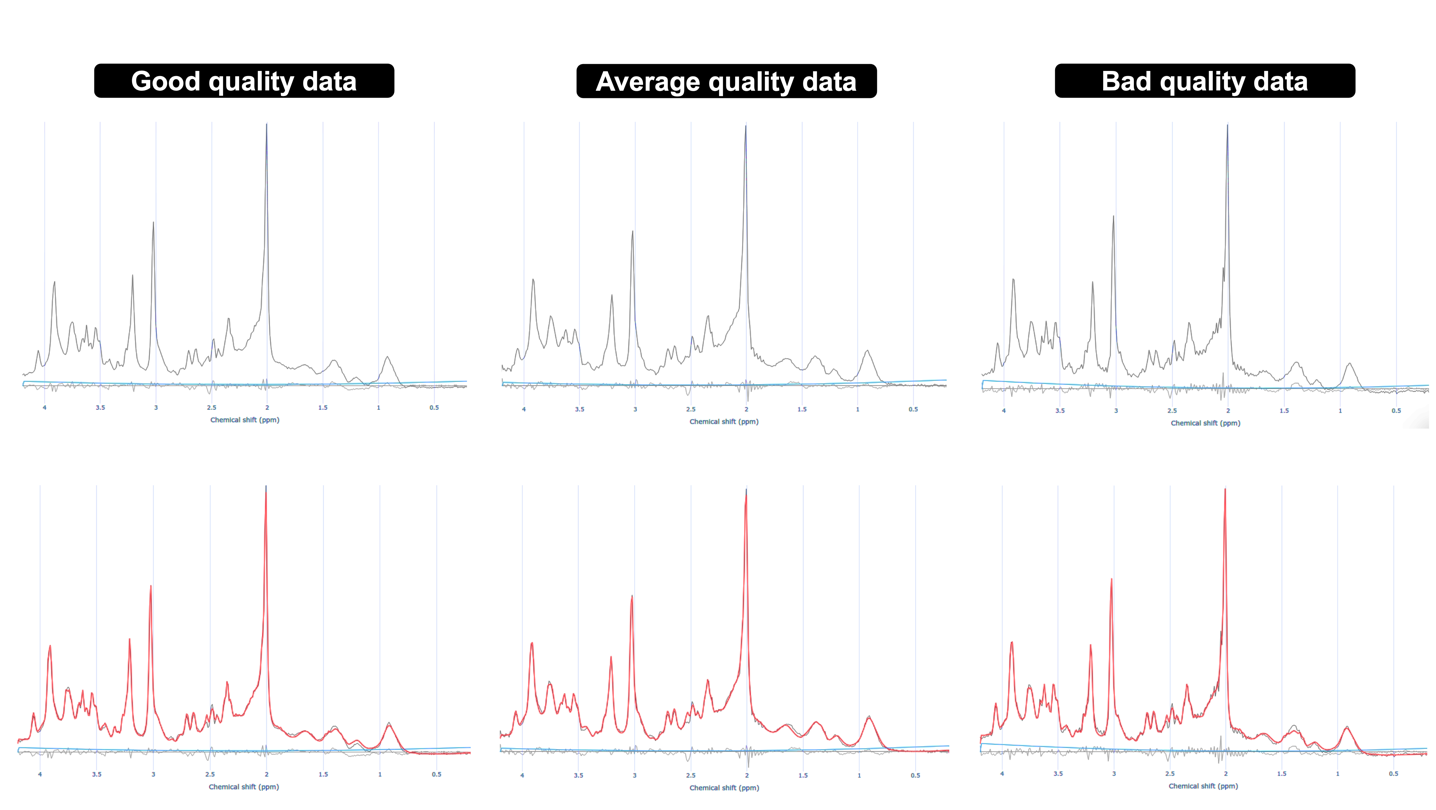

**Figure S1. Representative spectra.** Representative spectra from the hand knob (motor cortex) acquited on a 7T scanner. Both raw (upper row) and fitted data (lower row) are shown. From left to right, good, average and bad data quality spectra are shown respectively. The chemical shift axis is labeled in ppm units.
